# Loss of cGMP-dependent protein kinase II alters ultrasonic vocalizations in mice, a model for speech impairment in human microdeletion 4q21 syndrome

**DOI:** 10.1101/2021.01.06.425531

**Authors:** Tiffany M. Tran, Jessica K. Sherwood, Matheus F. Sathler, Franz Hofmann, Leslie M. Stone-Roy, Seonil Kim

## Abstract

Chromosome 4q21 microdeletion leads to a human syndrome that exhibits restricted growth, facial dysmorphisms, mental retardation, and absent or delayed speech. One of the key genes in the affected region of the chromosome is *PRKG2*, which encodes cGMP-dependent protein kinase II (cGKII). Mice lacking cGKII exhibit restricted growth and deficits in learning and memory, as seen in the human syndrome. However, speech/vocalization impairments in these mice have not been determined. Moreover, the molecular pathway underlying speech impairment in humans is not fully understood. Here, we employed cGKII knockout (KO) mice as a model for the human microdeletion syndrome to test whether vocalizations are affected by loss of the *PRKG2* gene. Mice emit ultrasonic vocalizations (USVs) to communicate in social situations, stress, and isolation. We thus recorded ultrasonic vocalizations as a model for speech in humans. We isolated postnatal day 5-7 pups from the nest to record and analyze USVs and found significant differences in vocalizations of KO mice relative to wild-type and heterozygous mutant mice. KO mice produced fewer calls that were shorter duration, higher frequency, and lower intensity. Because neuronal activity in the hypothalamus is important for the production of animal USVs following isolation from the nest, we assessed hypothalamic activity in KO pups following isolation. Indeed, we found abnormal hyperactivation of hypothalamic neurons in cGKII KO pups after isolation. Taken together, our studies indicate that cGKII is important for neuronal activation in the hypothalamus, which is required for the production of USVs in neonatal mice. We further suggest cGKII KO mice can be a valuable animal model for human microdeletion 4q21 syndrome.

**Highlights:**

- Chromosome 4q21 microdeletion leads to a human syndrome that exhibits restricted growth, mental retardation, and absent or delayed speech.
- The *cGMP-dependent protein kinase II (cGKII)* gene is one of the genes located in the affected region of the chromosome.
- cGKII knockout mice show restricted growth and deficits in learning and memory.
- Altered ultrasonic vocalizations and abnormal activation in hypothalamic neurons are found when infant cGKII knockout pups are isolated from the nest.
- cGKII knockout mice can be a valuable animal model for human microdeletion 4q21 syndrome.

## Introduction

Microdeletions of chromosome 4q21 are associated with a human syndrome that includes restricted growth, mental retardation, facial dysmorphisms, and absent or delayed speech [1, 2]. The affected region contains five genes; *PRotein Kinase G 2 (PRKG2), RasGEF Domain Family Member 1B (RASGEF1B), Heterogeneous Nuclear Ribonucleoprotein D (HNRNPD), Heterogeneous nuclear ribonucleoprotein D-like (HNRPDL)* and *Enolase-Phosphatase 1 (ENOPH1)* [1, 2]. Studies of human patients with different but overlapping deletions suggest that disruption of the *PRKG2* and *RASGEF1B* genes are important for intellectual disability and speech defects, while defects in *HNRNPD* and *HNRNPDL* genes are involved in growth retardation, hypotonia, and some aspects of developmental delay [2]. However, the exact molecular pathway underlying the symptoms in the human microdeletion syndrome has not been fully elucidated.

Among genes located in the affected chromosome region, *PRKG2* encodes cGMP-dependent protein kinase II (cGKII), which is a serine/threonine kinase activated by the second messenger cGMP [3, 4]. It is expressed abundantly in several tissues including the intestines, kidney, and brain and found at lower levels in the lungs, prostate, growth plates, pancreas and salivary glands [4, 5]. cGKII phosphorylates various targets that are important for several biological functions. For example, it plays a key role in modulating secretion from cells in the kidney, epithelium, and adrenal gland by phosphorylating the Cystic Fibrosis Transmembrane Conductance Regulator (CFTR) chloride channel [3, 5-8]. Moreover, loss-of-function mutations or genetic ablation of cGKII cause dwarfism in humans and rodents as cGKII regulates bone mass [6, 9–11]. In neurons, cGKII plays important roles in fast synaptic transmission by controlling the AMPA receptor (AMPAR) subunit GluA1 trafficking [12–14]. In particular, phosphorylation of serine 845 of GluA1 (pGluA1) is important for activity-dependent trafficking of GluA1-containing AMPARs and increases the level of extrasynaptic receptors [12, 15–18]. In the hippocampus, cGKII-mediated GluA1 phosphorylation is critical for long-term potentiation (LTP), which is important for learning and memory [12, 13, 19]. In fact, cGKII knockout (KO) mice exhibit severe impairment in hippocampus-dependent learning and memory [20, 21], which is relevant to mental retardation found in human microdeletion 4q21 syndrome [1, 2]. Taken together, loss of the *cGKII* gene may underlie neurological symptoms in microdeletion 4q21 syndrome, which includes intellectual disability and speech defects.

Verbal language is specific for humans and some species, including non-human primates, but most animals largely lack verbal vocalization as a communication tool [22]. Speech production involves multiple brain regions as well as neuronal control of muscles in the lung, larynx, and pharynx [23–25]. Significantly, speech defects are one of the primary characteristics of the human microdeletion 4q21 syndrome, but the mechanisms underlying speech delay are unknown [1,2]. Rodents communicate through the use of high frequency, ultrasonic vocalizations (USVs), and the specific frequency emitted is dependent on the context, including male-female interactions, juvenile social interactions, and in mother-infant interactions [22, 26-28]. Pups vocalize in the range of 30-90 kHz in response to aversive stimuli, such as isolation from the natal nest, and this elicits a retrieval response from the mother [29–34]. Therefore, these calls have been thought to function as communication between animals [34]. Interestingly, isolation from the nest results in a decrease in body temperature of pups, which in turn triggers emission of USVs [31]. Notably, a recent study demonstrated that dropping body temperature of isolated pups induces strong neuronal activation in the hypothalamus following 90 min isolation from the nest, which is sufficient to produce USVs [35]. Therefore, thermal insulation is the important factor regulating the activation of hypothalamic neurons in pups following isolation from the nest to emit USVs [35].

Given that cGKII KO mice exhibit dwarfism and disrupted learning and memory, which resembles traits of human patients with microdeletion 4q21 syndrome, these mice can be a potential animal model to study this disorder. However, whether defects in the *cGKII* gene are sufficient to produce vocalization changes in addition to the cognitive defects and growth retardation seen in KO animals is unknown. Here, we measure USVs to examine whether loss of the cGKII gene underlies speech impairment in human syndrome. We found significant differences in USVs of KO mice relative to wild-type (WT) mice. Furthermore, we reveal abnormal activation in cGKII KO neurons in the hypothalamus that play important roles in the production of USVs.

Our studies thus demonstrate that cGKII is important for USVs in mice. Taken together, the current study combined with previous work demonstrating the effects due to loss of cGKII function on learning, memory and bone growth provide strong evidence that cGKII is the valuable animal model for human microdeletion 4q21 syndrome.

## Materials and Methods

### Animals

cGKII KO animals were maintained in the C57Bl6 background as previously described [20]. Pups, a mixture of males and females, were produced from breeding heterozygous cGKII KO mice. Litters consisting of a mixture of WT, KO, and heterozygous mutant mice were used for the study. Colorado State University’s Institutional Animal Care and Use Committee reviewed and approved the animal care and protocol (16-6779A).

### USV Recording and analysis

A decrease in body temperature of pups upon isolation from the nest triggers emission of USVs. However, after the second week of life, pups are able to self-maintain their body temperature, resulting in fewer USVs [31]. Therefore, the ages of the pups used for experiments ranged from postnatal day 5 to postnatal day 7 (P5-P7) as reported previously [36]. Neonatal P5-7 mice were separated from the home cage and placed in a round transparent container with an open top. The ultrasound microphone (Avisoft-Bioacoustics CM16/CMPA) was placed above the container in the sound-proof box. Vocalizations were recorded for 5 min using Avisoft Recorder software. After the recording was finished for each pup, tail biopsy was carried out for genotyping, and brain tissues were collected for analyzing cGKII expression. Between animals, the container was cleaned with 70% ethanol and dried to get rid of any odors that could affect pup vocalizations. Spectrograms of USVs were generated and analyzed by using Avisoft SASLab pro (**Fig. 1**). Recording data were filtered in a range of 30-90kHz, and start/end time, duration, peak frequency, and peak amplitude for each of the calls were collected. With the data extracted, the number of calls, average duration, average frequencies, and average amplitudes were determined for each pup individually and then averaged for each genotype.

**Figure 1.**
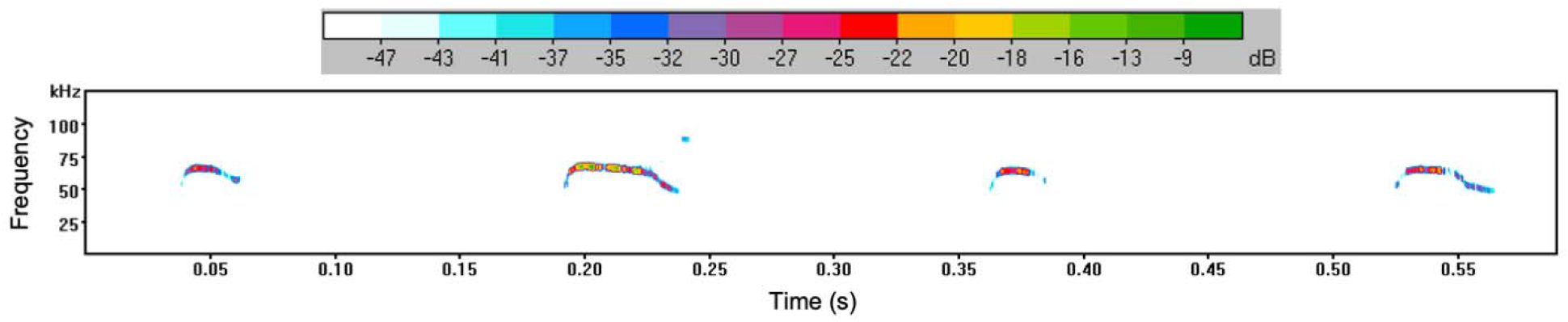
Spectrogram of mouse pup ultrasonic vocalizations. Spectrogram generated from P5-P7 pup isolation calls using Avisoft SASLab Pro software. Four individual vocalizations are shown in the representative spectrogram. The horizontal axis shows time (in seconds) from which duration of each USV could be determined as well as a total duration of vocalization during the 5-minute testing period. The vertical axis shows frequency (kHZ) of vocalizations emitted by mouse pups. The amplitude of calls (dB) can also be determined by the color (scale shown in top bar).

### Tissue sample preparation and immunoblots

Whole brain tissue from mouse brain was prepared as described previously [20, 37–39]. Brain samples were homogenized and collected by using a RIPA buffer (150mM NaCl, 0.1% TritonX-100, 0.5% sodium deoxycholate, 0.1% SDS, and 50 mM Tris, pH 8.0). Equal amounts of protein were loaded on 10% SDS-PAGE gel and transferred to nitrocellulose membranes. Membranes were blotted with anti-cGKII [12, 20] (Covance, 1:1000) and anti-actin (Abcam, 1:2000) antibodies and developed with chemiluminescence.

### Immunohistochemistry

Immunohistochemistry was carried out as described previously [37]. 5-day old WT and cGKII KO pups (both males and females) were subjected to 90 mins isolation from the nest. Both WT and KO pups in the nest were used as the basal condition. Brains were then extracted and fixed with 4% paraformaldehyde for 3 days, and then 40μm coronal sections were collected using a vibratome. The expression of the activity-regulated gene, c-Fos, was used to determine activation of hypothalamic neurons before or after isolation from the nest. Sections containing the hypothalamus were collected, permeabilized with diluting buffer (1% BSA, 0.4% Triton-X, 1% normal goat serum in pH 7.6 TBS), and blocked with 3% normal goat serum in TBS. The brain tissues were then incubated with anti-c-Fos antibody (Synaptic Systems) diluted to 1:500 in diluting buffer at 4°C for 18 hours. Afterwards, the sections were incubated for 2 hours with a secondary goat anti-rabbit IgG antibody conjugated with Alexa Fluor 647 (Invitrogen). The sections were then coverslipped and mounted on microscope slides. Sections were imaged using the Olympus inverted microscope IX73. The Olympus cellSENS Dimensions software was used to measure the area fraction of c-Fos positive immunoreactivity in the hypothalamic region.

### Statistics

Statistical comparisons were analyzed with the GraphPad Prism 8 software. Unpaired two-tailed Student t-tests were used in single comparisons. For multiple comparisons, a oneway ANOVA followed by a Fisher’s LSD test was used to determine statistical significance. Results are represented as mean ± SEM, and *p* < 0.05 was considered statistically significant.

## Results

### Loss of cGKII function in mouse pups disrupts USVs

Following isolation from the nest, we recorded USVs from P5-7 pups of WT, cGKII KO, and heterozygous mutant mice. As mouse pup USVs occur in the 40-80 kHz range [40], spectrograms in this range for each recording were created and analyzed (**Fig. 1**). We found that pups lacking the *cGKII* gene emitted less vocalizations than both WT and heterozygous mutant (HET) pups during the 5 min testing period (WT, 397.9 ± 37.55 calls; KO, 251.4 ± 29.51 calls; HET, 401.4 ± 28.52 calls; p=0.0067) (**Fig. 2a**). The duration of each call of KO pups was also significantly reduced compared with WT and HET mice (WT, 79.57 ± 7.51 ms; KO, 50.25 ± 5.90 ms; HET, 80.28 ± 5.70 ms; p=0.0067) (**Fig. 2b**). We also analyzed the peak frequency of USVs and found that KO pups produced higher frequency USVs than WT and HET pups (WT, 64.85 ± 0.60 kHz; KO, 67.51 ± 1.01 kHz; HET, 64.6 ± 0.53 kHz; p=0.0133) (**Fig. 2c**). When comparing the peak amplitude of USVs from each genotype, USVs from WT pups had a larger amplitude than from KO pups (WT, −15.76 ± 1.03 dB; KO, −18.62 ± 1.27 dB; HET, −16.15 ± 0.52 dB; p=0.04) (**Fig. 2d**). Given that KO pups generated less and shorter calls (**Fig. 2a and 2b**), the total time of USVs for 5 min in KO was significantly less than WT and HET mice (WT, 35.32 ± 5.88 s; KO, 14.56 ± 2.58 s; HET, 36.29 ± 4.58 s; p=0.0106) (**Fig. 2e**). Interestingly, there were no significant differences between WT and HET pups. Taken together, these results show that the complete loss of the *cGKII* gene in mice is sufficient to impair USVs.

**Figure 2.**
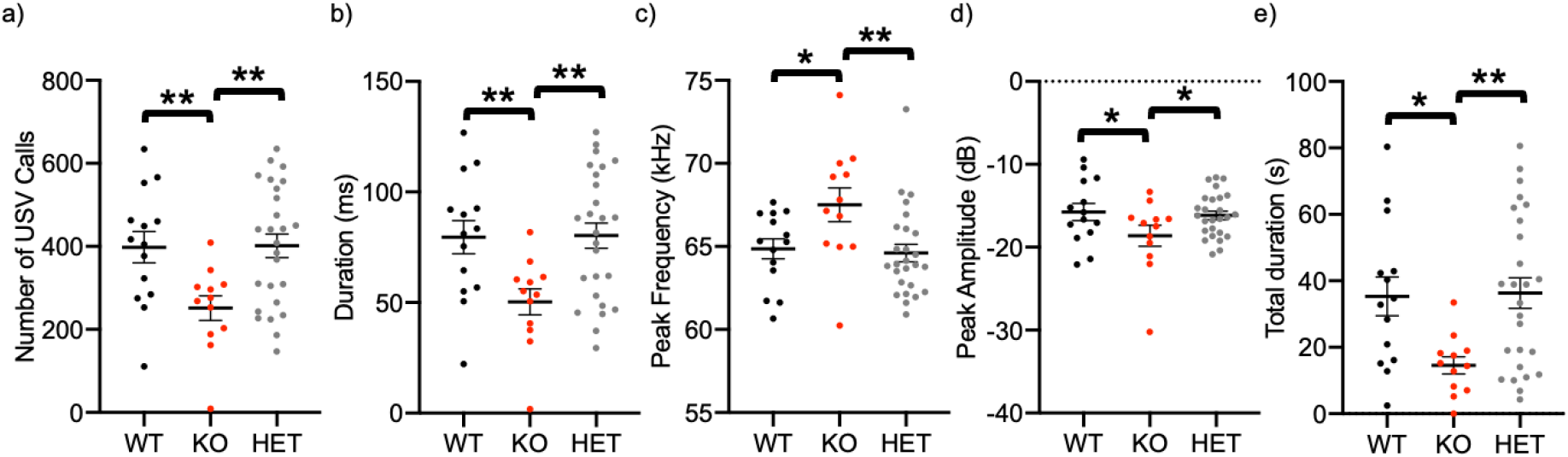
Abnormal ultrasonic vocalizations in isolated cGKII KO pups relative to WT and HET pups. **a)** Total number of USVs emitted, **b)** duration of individual calls, **c)** peak frequency, **d)** peak amplitude (or intensity), and **e)** total duration of calls in each genotype during 5-minute isolation period. (n=14 WT, 12 KO, and 26 HET animals, *p<0.05 and **p<0.01, one-way ANOVA, Fisher’s LSD test).

### WT and HET mice exhibit normal cGKII expression

Interestingly, for all characteristics of USVs, such as number of calls, duration of each call, peak frequency, and peak amplitude, there were no significant differences between WT and HET pups (**Fig. 2**). To determine whether cGKII protein is lower in HET pups relative to WT pups, we isolated brain tissues from each genotype and measured cGKII expression levels by immunoblots (**Fig. 3**). As expected, cGKII KO animals had no cGKII expression, whereas there was no difference in cGKII levels between WT and HET brains (WT, 1 ± 0.10 a.u. and HET, 1.07 ± 0.15 a.u.) (**Fig. 3**). Therefore, one copy of the *cGKII* gene in mice is sufficient to produce sufficient cGKII protein, ultimately leading to normal USV phenotypes in HET pups.

**Figure 3.**
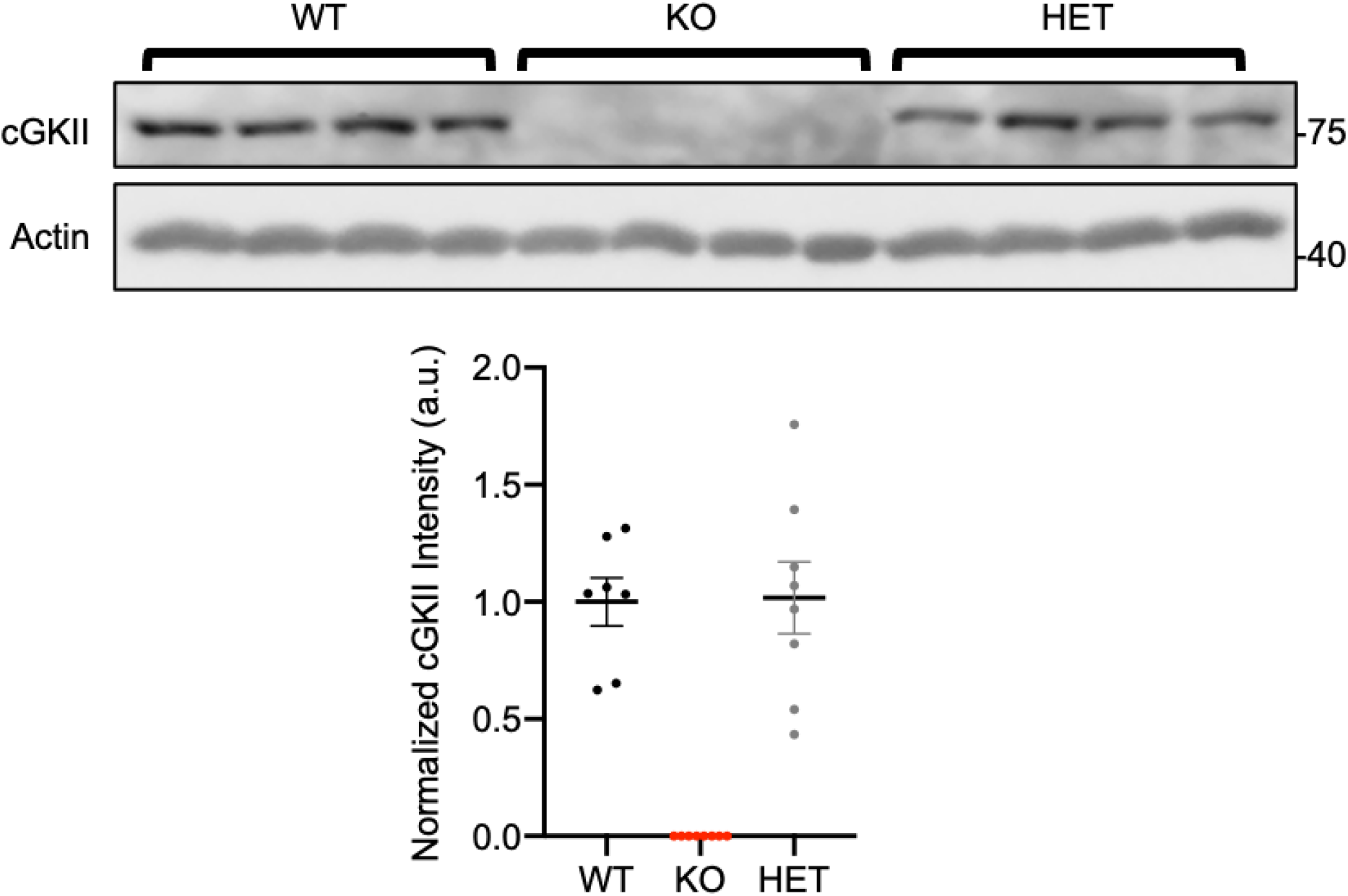
cGKII protein expression levels in each genotype. Representative immunoblots of cGKII protein expression in brain tissues from 4 animals in each genotype. Actin protein expression is used as a loading control. Normalized cGKII protein levels are shown in graph below. KO pups have no protein expression, while WT and HET pups have the same amount of protein present.

### Abnormal activation of cGKII KO neurons in the hypothalamus

Given that the activity of neurons in the hypothalamus is important for USV emission in infant mice [35] and cGKII is able to regulate neuronal activity by controlling AMPAR trafficking [12–14], we hypothesized that loss of cGKII function can alter neuronal activation in the hypothalamus following isolation from the nest, which in turn affects USVs. We thus measured expression of the activity-regulated gene, c-Fos, as a neuronal activity marker [41,42] to determine activation of hypothalamic neurons before or after isolation (**Fig. 4 and Supplementary Fig. 1**). WT mice after a 90 min isolation exhibited significantly increased neuronal activity in the hypothalamus compared to WT mice in the nest (WT nest, 0.45 ± 0.08 % and WT isolation, 0.86 ± 0.15 %, p=0.0411) (**Fig. 4**), which is consistent with the previous report [35]. There was no difference in c-Fos immunoreactivity of WT and KO mice before the isolation (WT nest, 0.45 ± 0.08 % and KO nest, 0.49 ± 0.11 %) (**Fig. 4**). Interestingly, a 90 min isolation significantly elevated c-Fos immunoreactivity in the hypothalamus of KO mice compared to KO nest pups (KO nest, 0.49 ± 0.11 % and KO isolation, 1.70 ± 0.30 %, p<0.0001) (**Fig. 4**). More importantly, increased neuronal activation in isolated KO animals was significantly higher than neuronal activity in WT pups following isolation (WT 0.86 ± 0.15 % and KO 1.70 ± 0.30 %, p=0.0005) (**Fig. 4**). These results thus suggest that the loss of cGKII is associated with an abnormal increase in the neuronal activity in the hypothalamus following isolation, which may underlie altered USVs in KO mice.

**Figure 4.**
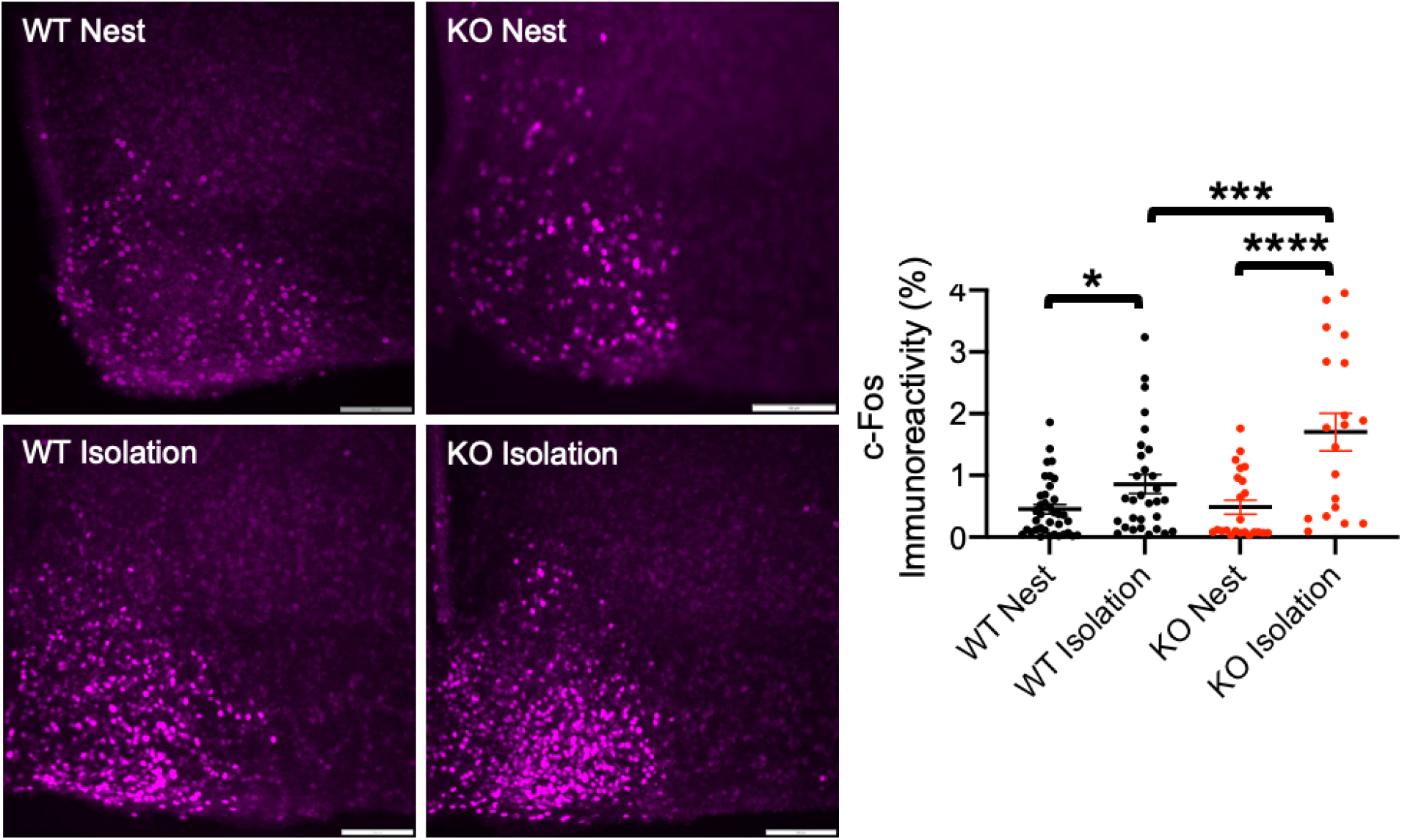
Hyperactivation of hypothalamic neurons in isolated cGKII KO pups. Left panels: Representative c-Fos immunohistochemistry images in each condition. The graph on the right shows that isolation from the nest elevates neuronal activity in the WT hypothalamus, but abnormal overactivation is found in isolated KO pups (n= WT Nest; 37 sections from 5 animals, WT Isolation; 30 sections from 4 animals, KO Nest; 23 sections from 3 animal, and KO Isolation; 19 sections from 2 animals, *p<0.05, ***p<0.001, and ****p<0.0001, one-way ANOVA, Fisher’s LSD test). A scale bar indicates 100μm.

## Discussion

4q21 microdeletion syndrome is a human genomic disorder, which is characterized by neonatal hypotonia, intellectual disability, absent or delayed speech, growth retardation, and brain malformation and facial dysmorphism [1, 2]. Given that mice carrying a null mutation of the *cGKII* gene exhibit postnatal dwarfism and learning disability similar to that observed in the reported patients [6, 20, 21], it has been proposed that the *PRKG2* gene is one of the key genes for the 4q21 microdeletion syndrome [1,2]. Additionally, cGKII is abundantly expressed in the brain and regulates neuronal activity and cognitive function [12, 13, 20, 21]. However, a role of cGKII in speech has not been investigated. Here, we recorded USVs in neonatal mice and revealed abnormal vocalization patterns in mice lacking the *cGKII* gene, suggesting cGKII plays a critical role in the production of USVs. Therefore, loss of cGKII function can be a valuable animal model to investigate pathophysiology of the 4q21 microdeletion syndrome.

A lack of cGKII expression in a specific area of the brain responsible for the production of vocalization in fetal life could be a possible reason for USV changes in mice. In fact, isolation from the nest decreases body temperature in neonatal mice, which in turn activates neurons that express agouti-related protein (AgRP) in the hypothalamus [35]. This activation is sufficient to produce USVs, so that the mother can retrieve isolated pups to the nest [29–34]. Given that cGKII is important for neuronal activity, loss of cGKII function would be expected to lower the activity of AgRP neurons in the hypothalamus, which may underlie abnormal USVs in KO animals. In contrast to our prediction, the neuronal activity measured by c-Fos immunoreactivity in KO mice was not significantly different from WT pups in the nest. (**Fig. 4**). This suggests cGKII may not be important for neuronal activity under the basal condition. Surprisingly, isolation from the nest induced overactivation in hypothalamic neurons (**Fig. 4**). Notably, hypothalamic AgRP neurons are regulated by GABAergic inhibitory interneurons [43], and this regulation requires the cold-sensitive transient receptor potential cation channel, TRPC5 [43, 44]. This suggests loss of cGKII function may decrease activity in cold-sensitive GABAergic neurons, which in turn abnormally elevates AgRP neuronal activity in the hypothalamus, altering USV production in KO pups. Nonetheless, we are unable to rule out the possibility that loss of the *cGKII* gene in other brain regions and non-neuronal tissues may be involved. In particular, the larynx has skeletal muscles required for USV generation in mice [22]. Given that cGKII is expressed in skeletal muscles [45], altered USVs in cGKII KO mice may be impacted by deficits in larynx muscles.

As infant mouse USVs after isolation trigger the mother to retrieve the isolated pups to the nest [34], the maternal retrieval time for KO pups may be longer than WT mice. This would examine whether USVs generated from isolated pups indeed are used for animal communication, similar to human speech. Therefore, further studies on USVs and retrieval assays of cGKII KO pups may provide helpful insight to human disorders associated with deficits in social communication ability in the 4q21 microdeletion syndrome.

## Acknowledgements

We thank members of the Kim laboratory for their generous support and Dr. Lon Kendall for use of the USV recording system. We also thank Dr. Ioana Carcea at Rutgers University for helping USV data analysis.

## Funding

This work is supported by Student Experiential Learning Grants from Colorado State University (LMS and SK), the pilot program from the Colorado Clinical and Translational Sciences Institute (SK), College Research Council Shared Research Program from Colorado State University (SK), and the Boettcher Foundation (SK).

---

**Supplementary figure 1.**
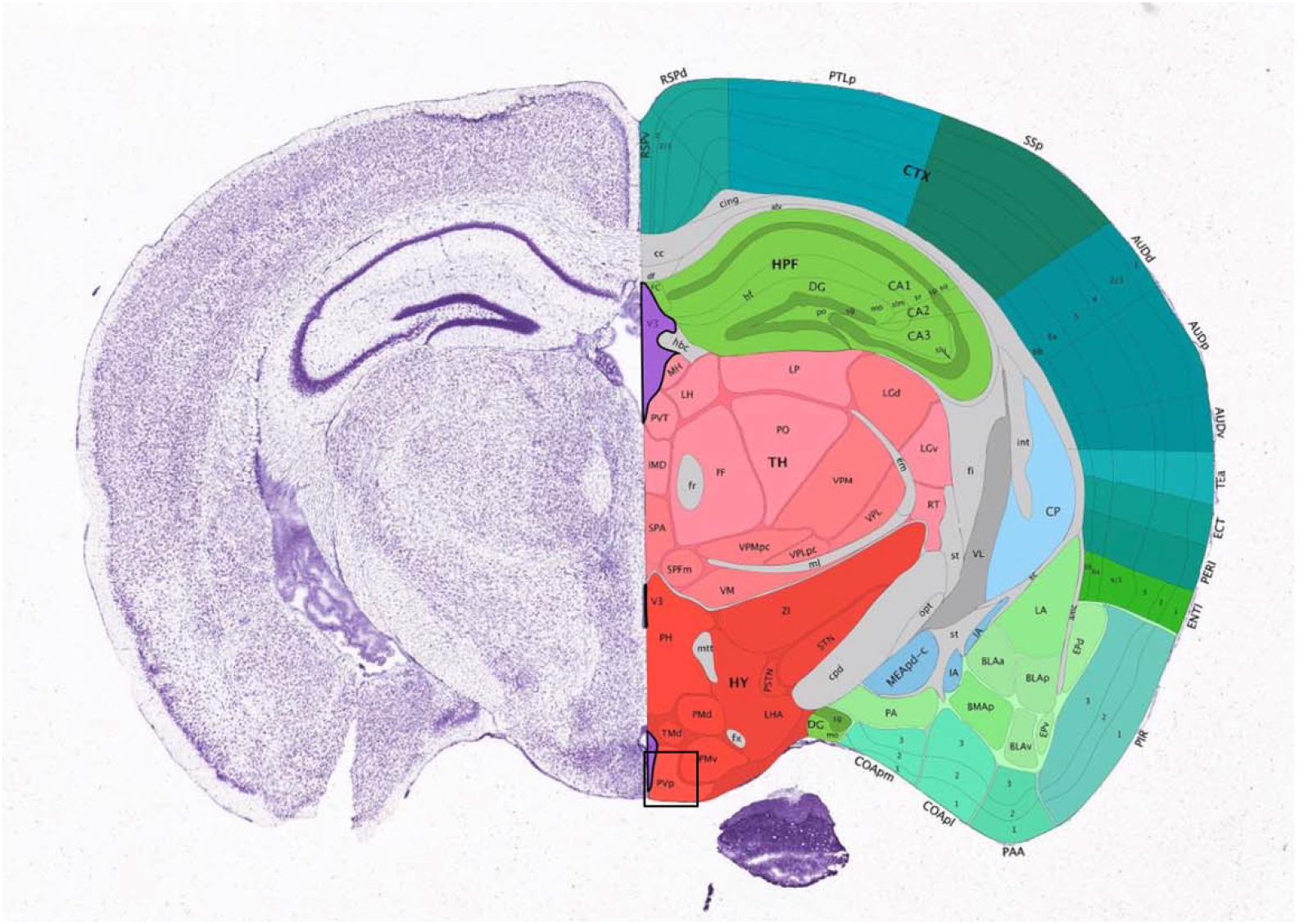
The mouse brain atlas from the Allen Brain Institute (https://mouse.brain-map.org/experiment/thumbnails/100048576?image_type=atlas) showing the sub-brain region (hypothalamus, outlined by the black square) used for quantification in Fig.

## Notes

### Competing Interest Statement

The authors have declared no competing interest.

